# Reduced foraging investment as an adaptation to patchy food sources: a phasic army ant simulation

**DOI:** 10.1101/101600

**Authors:** Serafino Teseo, Francesco Delloro

## Abstract

Colonies of several ant species within the subfamily Dorylinae alternate stereotypical discrete phases of foraging and reproduction. Such phasic cycles are thought to be adaptive because they minimize the amount of foraging and the related costs, and at the same time enhance the colony-level ability to rely on patchily distributed food sources. In order to investigate these hypotheses, we use here a simple computational approach to study the population dynamics of two species of virtual ant colonies that differ quantitatively in their foraging investment. One species, which we refer to as “phasic”, forages only half of the time, mirroring the phasic activity of some army ants; the other “non-phasic” species forages instead all the time. We show that, when foraging costs are relatively high, populations of phasic colonies grow on average faster than non-phasic populations, outcompeting them in mixed populations. Interestingly, such tendency becomes more consistent as food becomes more difficult to find but locally abundant. According to our results, reducing the foraging investment, for example by adopting a phasic lifestyle, can result in a reproductive advantage, but only in specific conditions. We thus suggest phasic colony cycles to have emerged together with the doryline specialization in feeding on the brood of other eusocial insects, a resource that is hard to obtain but highly abundant if available.

## Introduction

Within several taxa belonging to the ant subfamily Dorylinae (*sensu* Brady et al., 2014), species are considered “phasic” or “non-phasic” according to their lifestyle (Schneirla, 1971; Gotwald, 1995; Kronauer, 2009). In phasic species, broods develop synchronously in distinct cohorts of the same age, and colonies undergo virtually discrete phases of foraging and reproduction based on the presence or absence of food-demanding larvae (Schneirla, 1971; Gotwald, 1995; Ravary and Jaisson, 2002; Kronauer, 2009; Teseo et al., 2013). In non-phasic species, larvae are not synchronized in their development, and foraging and reproduction are not strictly coordinated with reproductive cycles. Within army ants, non-phasic taxa include the genera *Dorylus* and *Labidus*, whereas phasic taxa include some species within the Old World genus *Aenictus* and the New World genera *Eciton* and *Neyvamyrmex* (Kronauer, 2009). Outside army ants, phasic groups include some species in the genera *Sphinctomyrmex* (Buschinger et al., 1989), *Leptanilloides* (Brandão et al., 1999; Donoso et al., 2006), *Cerapachys* (Wilson, 1958a; Ravary and Jaisson, 2002), and to some extent *Simopelta* within the Ponerinae subfamily (Gotwald and Brown, 1967), and *Leptanilla japonica* within Leptanillinae (Masuko, 1990). Found in loosely related groups, phasic colony cycles have recently been suggested to have evolved repeatedly and early in army ant evolution, and to have been secondarily lost in genera such as *Dorylus* and *Labidus* (Kronauer, 2009).

Whereas the phasic cycles of *Eciton* and *Neyvamyrmex* have been extensively studied in the field throughout the 20th century (Hagan, 1954a, 1954b, 1954c, Schneirla, 1934, 1945, 1944a, 1944b; Topoff et al., 1980; Topoff, 1984), the “clonal raider ant” *Cerapachys biroi* is the only species in which the mechanistic aspects of phasic colony cycles have been thoroughly studied in highly controlled laboratory experiments (Ravary and Jaisson, 2002; Ravary et al., 2006; Teseo et al., 2013; Ulrich et al., 2015). In this parthenogenetic queenless species, the presence of larvae inhibits the ovarian activation in workers and stimulates foraging behavior. This results in developmentally synchronized cohorts of larvae and phasic foraging activity limited to when larvae are present.

Although the molecular, individual and colony-level mechanisms underlying the alternation of phases are now beginning to be understood (Ravary and Jaisson, 2002; Ravary et al., 2006; Teseo et al., 2013; Oxley et al., 2014; Ulrich et al., 2015), the adaptive significance of the phasic lifestyle is still to some extent unknown. At present, a single study (Kronauer, 2009) has formulated explicit hypotheses about the adaptive value of the phasic lifestyle, suggesting that it likely provides several main benefits. First, in some army ant species, colonies migrate throughout the foraging phases. Migrations probably maximize the foraging success and help avoiding resource depletion at a local scale (Wilson, 1958b; Franks and Fletcher, 1983; Gotwald, 1995). Second, phasic cycles minimize the time invested in foraging, which in turn minimizes the costs involved in foraging activity and emigrations. Third, stationary reproductive phases are sometimes necessary because, in raid-conducting genera such as *Eciton* and *Neyvamyrmex* (but not in *C. biroi*), physogastric egg-laying queens are not mobile or cannot be transported by workers. Finally, doryline ants are in fact specialized predators of the brood of other social insect colonies (Brady et al., 2014; Borowiec, 2016), a food source that is difficult to find but may be overabundant when found (Kronauer, 2009). Developmentally synchronized cohorts of larvae should, in principle, consume more efficiently the large quantities of rapidly decaying prey that become unpredictably available during foraging phases compared to non-synchronized broods including eggs and pupae.

In this study, we use simple computer simulations to explore some of the hypotheses regarding the adaptive value of phasic cycles in army ants, with the goal of understanding whether these may have appeared as an adaptation to specific ecological conditions. In nature, colonies of phasic ant species only forage in the presence of developing larvae, a period corresponding approximately to half the duration of a complete reproductive cycle in *E. burchelli* or *C. biroi* (Schneirla, 1971; Ravary and Jaisson, 2002). In such species, however, colonies need to reach a certain threshold size in order to successfully split, via fission, into two viable daughter colonies. Accordingly, the main assumption of our model is that, everything else being equal, phasic colonies invest in foraging only half of their time. Therefore, due to the reduced investment in foraging, growth and reproduction in phasic colonies are restricted to around only half of their potential. From an evolutionary perspective, in a hypothetical ancestral population in which different ant colonies make quantitatively different foraging investments, colonies with a low foraging investment should grow and reproduce relatively slower and eventually become extinct; on the other hand, colonies with a high foraging investment should grow and reproduce faster, invading the population. Our model aims to understand the evolution of phasic cycles by investigating the conditions in which reducing rather than maximizing the time invested in foraging may result in a selective advantage for virtual ant colonies. In particular, we ask whether and how the cost of foraging, the probability to find food items and the size of the food items affect the population dynamics of low- and high-foraging virtual ant colonies, which we respectively refer to as “phasic” and “non-phasic”. These two colony types, or species, behave exactly the same way, with the only exception that the foraging investment of phasic colonies is only half of that of non-phasic colonies.

In our simulations, we first examine the growth of monospecific populations of phasic or non-phasic colonies. Then, we explore the outcome of competition for food in mixed populations consisting of phasic and non-phasic colonies. We show that phasic colonies reproduce more than, and outcompete, non-phasic colonies when the cost of foraging is relatively high. Interestingly, this tendency becomes more consistent as food becomes more patchily distributed. Our results suggest that the more locally abundant and rare food sources are, the better phasic colonies outperform non-phasic colonies. Minimizing the foraging investment, for example by adopting a phasic lifestyle, could thus result in a reproductive advantage in specific conditions.

## The models

### Logistic density-dependent growth of phasic or non-phasic colonies

Our simulations are based on a variation of the continuous density-dependent growth model in single-species populations. In our model, growth depends on the food income and the cost of foraging. The growth of a colony can thus be expressed by the differential equation:

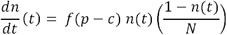

where *n(t)* is the number of individuals in the colony, *p* is the food income for the colony, *c* is the cost of foraging, *f* is a factor that describes the foraging investment and takes different values for phasic (*f*=1/2) and non-phasic (*f*=1) populations, and *N* is the maximal size that a colony can reach, expressed as its number of individuals. When *p=c*, the population growth equals zero and the system is in an unstable state of equilibrium. With p<c, the colony experiences negative growth and goes through extinction, whereas with p>c it grows until the maximal size is reached.

In our first set of simulations, we examine the growth curve of populations constituted of either only phasic or only non-phasic colonies, in discretized time. In each simulation, the total population is composed by a variable number of colonies and is represented by a vector 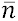, which we refer to as the population vector. Each element *n*_*i*_ within 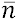 represents the number of individuals within the i-th colony. The length of 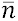 represents the number of colonies in the population, and varies with time depending on colony death and reproduction. At each time iteration (identified by the subscript *t*), the discrete variation of the size of each colony is computed according to the following equation:

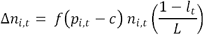

where *n*_*i,t*_ is the number of individuals in the i-th colony at the t-th time iteration, *f* is the “foraging” factor varying for phasic (*f*=1/2) and non-phasic (*f*=1) populations, *p*_*i,t*_ is the food income, *c* is the foraging cost, *l*_*t*_ is the number of colonies at time *t*, and *L* is the carrying capacity for the population, expressed as the maximal number of colonies that are able to survive in the environment. The idea behind our simulations is that virtual ant colonies explore the environment in search of food, and may encounter food items of various sizes and at different probabilities. The time unit of the simulation corresponds to one colony cycle, a period in which each colony may find a food item and, depending on its size, possibly reproduce or die. In our algorithm, colonies find on average the same quantity of food, but food distribution is parametrically controlled to allow testing and comparing differential scenarios. To keep the average food income constant among scenarios, the size of the available food items and the probability to encounter them are inversely proportional. For example, in a given scenario, food items are small and easy to find, whereas in another one they are large and difficult to find. We thus implement the food income *p*_*i,t*_ as a stochastic variable depending on the random number *x*, which is uniformly distributed in the interval [0,1], in the following way:

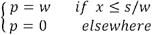

where *s* represents the average food income per colony, *w* is the parameter used to tune food distribution in the different scenarios (it regulates the probability of finding food without changing the average food income over time per colony), and *s/w* is the probability of finding food. The model is based on the iteration of an algorithm modelling a virtual ant colony cycle, which is taken as the time unit (Figure 1a). The algorithm begins with the generation of a random number *x*.. If *x* is smaller than *s/w*, the colony finds and consumes a quantity *w* of food, which results in a size increase; if *x* is larger than *s/w*, the colony does not encounter food and does not increase in size. In addition, as foraging is costly, colonies lose individuals at each time iteration. This limits the growth of the colonies that encounter food items, and results in negative growth for colonies that do not encounter food items. Changing the value of *w* modifies the probability at which each colony receives food and the size of the food items, but does not affect the average quantity of food received. For example, with *s* fixed at 0.4, if *w*=1 colonies receive food items of size 1 with a probability 0.4; on the other hand, if *w*=2, colonies receive food items of size 2 with a probability 0.2, and if w=0.5, they receive food items of size 0.5 with a probability 0.8 (Figure 1b). Implementing different values of *w* allows testing and comparing the performances of phasic and non-phasic colonies in various scenarios.

**Figure 1.**
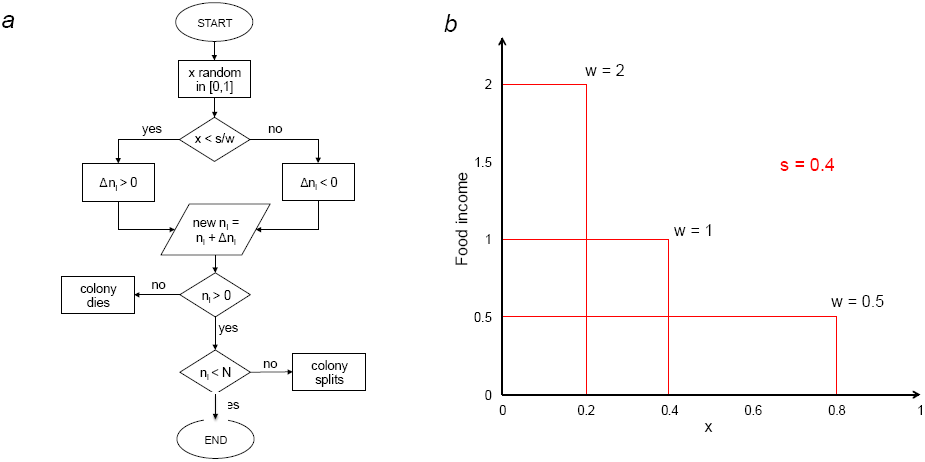
The simulation algorithm. ***a)*** At each time iteration, for each colony, our algorithm extracts a random number *x*. If *x* is smaller than *s/w*, the colony consumes a quantity *w* of food; if the value of the extracted *x* is larger than *s/w*, the colony does not consume any food, and does not grow in size. If a colony reaches the maximal size, it splits in two daughter colonies, whereas if its size falls below a given minimal value (one tenth of the maximal size), it dies. ***b)*** With *s* fixed at 0.4, if *w*=1 colonies receive food items of size 1 with a probability 0.4; if *w*=2, colonies receive food items of size 2 with a probability 0.2; if w=0.5, colonies receive food items of size 0.5 with a probability 0.8. The area underlying each of the red graphs is constant across cases.

Colonies of phasic army ant species generally reproduce via fission by splitting in two equally sized daughter colonies when they reach a certain threshold size. In order to reproduce such dynamics, the algorithm checks the size of each colony at the end of each time iteration. If a colony reaches the maximal size, which we refer to as *m*, it splits into two colonies of size *m*/2. If during the simulations the size of a given colony falls below the value *m*/10, the colony is considered not viable anymore, and is eliminated from the population vector. At the beginning of the simulations, the population always consists of a single colony of size *m*/2.

In the first set of simulations, we studied the dynamics of monospecific populations of phasic or non-phasic species, without inter-specific interactions. For each species, for each of 4 values of *w* (0.7; 1; 2; 4), we explored values of *c* uniformly distributed in the interval [0, 0.7]. We focused exclusively on the cases in which *p*≥*c* because populations always go through extinction when *p*<*c*. The duration of each simulation was set to 500 time units to allow populations to reach, in most cases, an almost stable state. Each simulation was repeated 1000 times. The maximal colony size (*m*) was fixed to 1000 individuals, and the carrying capacity *L* to 300 colonies. These values were changed to check their effect on the simulation outcome (data not shown), but as long as they were kept sufficiently high to assure a good statistical representation, they did not appear to affect the results.

### Competition between phasic and non-phasic colonies

Simulating the density-dependent growth of phasic or non-phasic colonies provides insight into the population dynamics of monospecific populations in various ecological conditions; however, it does not allow predicting the outcome of competition for food in populations made up of colonies of both species. In our second set of simulations, we implement a resource competition scenario within a mixed population of phasic and non-phasic colonies. In this case there are two population vectors, one for phasic colonies and the other for non-phasic ones, but the carrying capacity is computed on the total population, i.e. the sum of phasic and non-phasic colonies. Consistently with our monospecific model, at each time iteration (identified by the subscript *t*) the discrete size variation of each colony is computed according to the following equations, for phasic and non-phasic populations respectively:

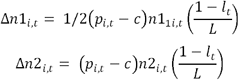

where *n*1 is the population vector of phasic colonies, *n*2 is the population vector of non-phasic colonies, *l*_*t*_ is the total number of colonies (phasic and non-phasic) at time *t*, and *L* is the carrying capacity expressed as the maximal number of colonies. Concerning reproduction, the same rules of our monospecific model apply here.

In this set of simulations, we studied the same values of *c*, *w* and s considered in the monospecific simulations. For each value of these parameters, we repeated the simulation 1000 times; each simulation consisted of 500 time iterations; we fixed *m*, the maximal colony size, to 1000, and the carrying capacity *L* to 300. Concerning the choice of such values, the same considerations made for the previous model apply here. All the simulations were carried out in a Python environment ({van Rossum}, 1995) with the module Numpy (Walt et al., 2011). Our scripts are provided in the supplementary electronic material.

## Results

### Logistic density-dependent growth of phasic and non-phasic colonies

Our results show that, as expected, both phasic and non-phasic colonies perform best when the cost of foraging is low. When the cost of foraging (*c*) exceeds a certain value that depends on *w* (the number that regulates the probability of finding food without changing the average quantity of food received over time by each colony), the population size at the end of the simulations decreases, and the probability of extinction increases (Figures 2a, b). When the cost of foraging and the probability to find food (*p*) approach the same value, i.e. close to the line described by the equivalence *p*=*c* (Figures 2a, b), the probability to go through extinction increases for both phasic and non-phasic populations. In general, at each value of *c*, the final population size increases with *p*, and the probability of extinction decreases (Figures 2a, b). The range of *c* and *p* values at which populations go through extinction becomes wider with increasing values of *w*. In particular, the extinction zone is wider for non-phasic colonies (Figure 2a) compared to phasic ones (Figure 2b), meaning that, for the same values of *c*, non-phasic colonies need more food than phasic colonies in order to avoid extinction.

**Figure 2.**
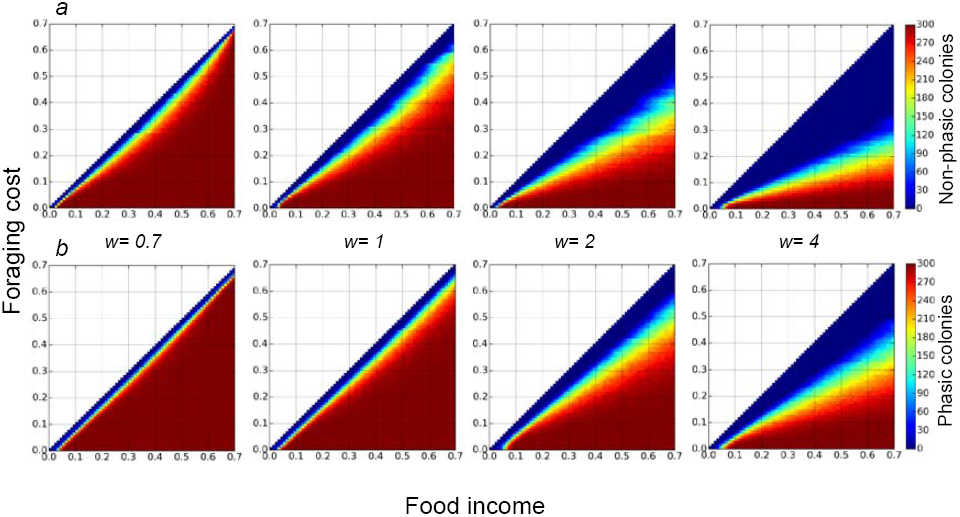
Monospecific populations of phasic and non-phasic colonies. ***a)*** The number of non-phasic colonies as a function of food income (*p* in the text) and foraging cost (*c* in the text), for four different values of *w*, in monospecific populations. ***b)*** Number of phasic colonies in monospecific populations. The color scale represents the mean number of colonies at the end of the simulations. As specified in the text, the probability of finding food decreases at increasing values of *w*, whereas the food item size increases.

For phasic colonies, for each value of *c* and at increasing values of *p*, the probability of extinction decreases and reaches zero (Figures 2b); on the other hand, for non-phasic colonies, the extinction probability decreases more slowly and never reaches zero, indicating that non-phasic colonies always face a risk of extinction (Figures 2a). Overall, higher extinction rates of non-phasic colonies lower their final population size across simulations.

### Competition between phasic and non-phasic colonies

At relatively low values of *w*, i.e. when food items are small but found frequently, non-phasic colonies outcompete phasic colonies and invade the population in most conditions (Figures 3a). On the other hand, phasic colonies tend to invade the population at values close to the line described by the equivalence *p=c*, in relatively more difficult conditions (Figures 3b).

As *c* increases, both populations need more food to avoid extinction, and with food becoming more difficult to find but locally abundant (i.e. at increasing values of *w*), survival becomes increasingly difficult for both types of colonies (Figure 3a). With w=4, for example, the carrying capacity is reached exclusively at relatively low foraging costs. However, similar to what happens in monospecific populations, phasic colonies go through extinction more slowly compared to non-phasic colonies, in that their prevalence increases at increasing values of *w* (Figure 3b). Even though for high values of *w* the total final population size is much lower than for low *w* values, phasic colonies outcompete non-phasic colonies in most conditions (Figures 3a, b).

**Figure 3.**
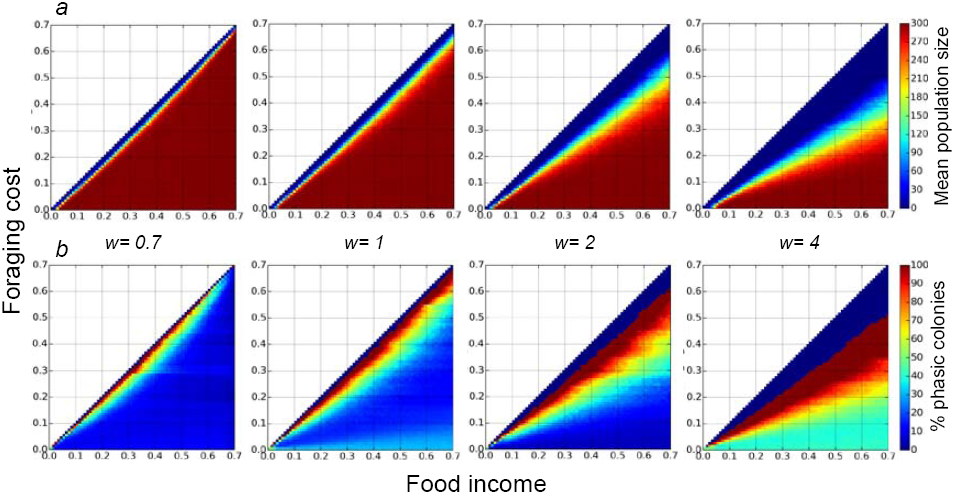
Mixed populations of phasic and non-phasic colonies. ***a)*** Mean size of mixed populations at the end of the simulations, as a function of food income (p in the text) and foraging cost (c in the text), for different values of w. As *w* increases, populations increasingly fail to reach the carrying capacity (300 colonies). ***b)*** The proportion of phasic colonies within mixed populations. The number of both phasic and non-phasic colonies decreases with the increase of w, whereas the prevalence of phasic colonies increases.

## Discussion

In this study, we have explored the population dynamics of two species of virtual ant colonies that differed quantitatively in their foraging investment. Our goal was to understand whether and when, under the assumptions of our model, colonies with a reduced foraging investment (or “phasic”) may have a better reproductive success compared to colonies that maximize their foraging effort (or “non-phasic”). We first studied monospecific populations of phasic or non-phasic colonies, and then mixed populations in which the two species competed for resources. When food was found frequently and the cost of foraging was set to relatively low values, monospecific populations of non-phasic colonies grew faster than phasic populations. On the other hand, due to their lower foraging investment, phasic colonies lost fewer individuals per iteration compared to non-phasic colonies, which resulted to be advantageous in challenging conditions. In fact, with the foraging cost set at relatively high values, phasic colonies survived through long series of iterations in which they could not access any food. This made them less likely to go through extinction, and also allowed them to reach larger population sizes compared to non-phasic colonies. Finally, and most interestingly, phasic populations performed increasingly better than non-phasic populations with food becoming rarer and locally abundant (higher values of *w*). Consistent to what we observed in monospecific populations, the dynamics of mixed populations depended as well on the distribution of the food items. When food was available only in small quantities but found easily, non-phasic colonies invaded the population in most conditions, and reached the carrying capacity of the environment. On the other hand, even though they did not reach the carrying capacity, phasic colonies tended to invade the population as food became rarer and more locally abundant, because they were less likely to go through extinction compared to non-phasic colonies.

According to our results, reducing rather than maximizing foraging activity can, in specific conditions, provide a significant reproductive advantage to ant colonies. Relying exclusively on costly, rare but locally abundant food is thus sustainable if the colony-level foraging investment is parsimonious; instead of foraging and raising brood continuously, phasic ants have evolved a system in which foraging, to which colony growth, reproduction and fitness are proportional, is reduced to phases virtually as short as the development of a single larva.

Most phasic ants, like many Dorylinae, are predators specialized in feeding on the brood of other social insects (Brady et al., 2014; Borowiec, 2016). Typically this is costly because prey colonies defend their nests and are in principle difficult to feed on. However, social insect colonies also generally house large amounts of brood, representing consistent food quantities for successful predator colonies. Our results thus suggest the possibility that the phasic lifestyle may have emerged together with the specialization in feeding on the brood of social insect colonies. Ecological pressures, such as inter-colony competition, might have pushed generalist non-phasic hunting ancestors to specialize in feeding on social insect brood. This might have selected for colonies that restricted their foraging investment, eventually resulting in the evolution of phasic cycles. In army ants, such a transition towards phasic activity may have occurred concurrently with the transition from individual to cooperative hunting (Wilson, 1958b; Hölldobler and Wilson, 1990; Kronauer, 2009), possibly through the re-activation of the gene networks responsible, in solitary-living Hymenoptera, for the alternation of foraging and reproductive phases (Amdam et al., 2006; Oxley et al., 2014).

The evolution of phasic activity cycles in ants needs to be further investigated a theoretical perspective. Our population dynamics approach, for example, could be integrated with a recently proposed mathematical model suggesting phasic colony cycles to be adaptive in species where the investment required to feed a single larva is high, but decreases as the number of larvae increases (Garnier and Kronauer, 2016). Previous research in theoretical ecology has made use of cellular automata to simulate the behavior of *Eciton burchelli* colonies (Britton et al., 1996), and analyze the effects on habitat quality and extension on population survival (Partridge et al., 1996; Britton et al., 1999). Using such cellular automata to model an interspecific competition scenario may help explore more specific questions about the adaptive value of phasic cycles. Such an approach would allow, for example, explicitly implementing the spatial and chronological dynamics of virtual ant colonies with differential degrees of phasic behavior, and testing whether and how phasic cycles may have a role in preventing colonies from depleting food resources at a local scale (Wilson, 1958b; Franks and Fletcher, 1983; Gotwald, 1995).

At present, empirically studying the adaptive value of phasic colony cycles appears to be challenging. A possible approach would likely consist in comparing phasic and non-phasic populations of the same or of closely related species. In particular, one could challenge such phasic and non-phasic colonies by feeding them either frequently with low amounts of food, or rarely with large amounts of food. Measuring the subsequent fitness of such experimental colonies would likely help understand whether phasic cycles are better suited for relying on rare but locally abundant food sources. At present, however, such an experiment would be hard to set up, mostly because phasic ant species are still poorly understood, cryptic and difficult to manipulate, especially in laboratory conditions. The only phasic ant that can be consistently studied in laboratory experiments is *C. biroi*. Using gene-silencing techniques on this species, an interesting approach would be “switching off” the mechanisms inhibiting worker ovarian activity in the presence of larvae, which would possibly result in producing non-phasic colonies. One could then test non-phasic and wild type colonies with differential feeding regimes.

The biology of doryline ants is still relatively poorly known. This is mainly because many taxa within the group are rare or difficult to encounter because of their hypogeaic lifestyle. In addition, a poor knowledge of the taxonomy of doryline ants, and a classification that until recently did not accurately take into account their evolutionary relationships have been an obstacle to comparative studies within the subfamily (Borowiec, 2016). A deeper understanding of the systematics and the behavior of Dorylinae is needed to gain new insights into the evolution of phasic colony cycles.

